# Interactive Tools for Functional Annotation of Bacterial Genomes

**DOI:** 10.1101/2024.04.15.589591

**Authors:** Morgan N. Price, Adam P. Arkin

**Affiliations:** Lawrence Berkeley National Lab

## Abstract

Automated annotations of protein function are error-prone because of our lack of knowledge of protein functions. For example, it is often impossible to predict the correct substrate for an enzyme or a transporter. Furthermore, much of the knowledge that we do have about the functions of proteins is missing from the underlying databases. We discuss how to use interactive tools to quickly find different kinds of information relevant to a protein’s function. Combining these tools often allows us to infer a protein’s function. Ideally, accurate annotations would allow us to predict a bacterium’s capabilities from its genome sequence, but in practice, this remains challenging. We describe interactive tools that infer potential capabilities from a genome sequence or that search a genome to find proteins that might perform a specific function of interest.

## Introduction

For most species of bacteria, we have genome sequences but not much experimental data about their capabilities. So, to understand these bacteria, we look at their genomes and try to predict phenotypes such as how they make energy, which carbon sources they can use, or which amino acids or vitamins they require for growth. In practice, this means predicting the protein-coding genes in the genome and then guessing at those proteins’ functions based on their similarity to proteins of known function (that is, whose function has been determined experimentally). This commentary describes how to use interactive tools to better predict the functions of proteins from diverse bacteria (and archaea) and the capabilities of those organisms.

Why are we focusing on interactive tools instead of fully automated annotations? We will show that automated annotations are intrinsically unreliable because much of our knowledge about proteins’ functions is missing from the underlying databases. In practice, automated annotations are often erroneous, even for proteins whose function is known. Automated annotations are necessary because interactive techniques require human attention and cannot be applied to every protein, but we view a protein’s automated annotation as a crude first guess. We focus on interactive tools -- ideally, web-based tools that run in a few seconds in a web browser -- because these make it easier to consider the wide range of additional data and analyses that can give insight into a protein’s function or a bacterium’s capabilities. An overview of our recommendations is shown in Figure 1.

**Figure 1:**
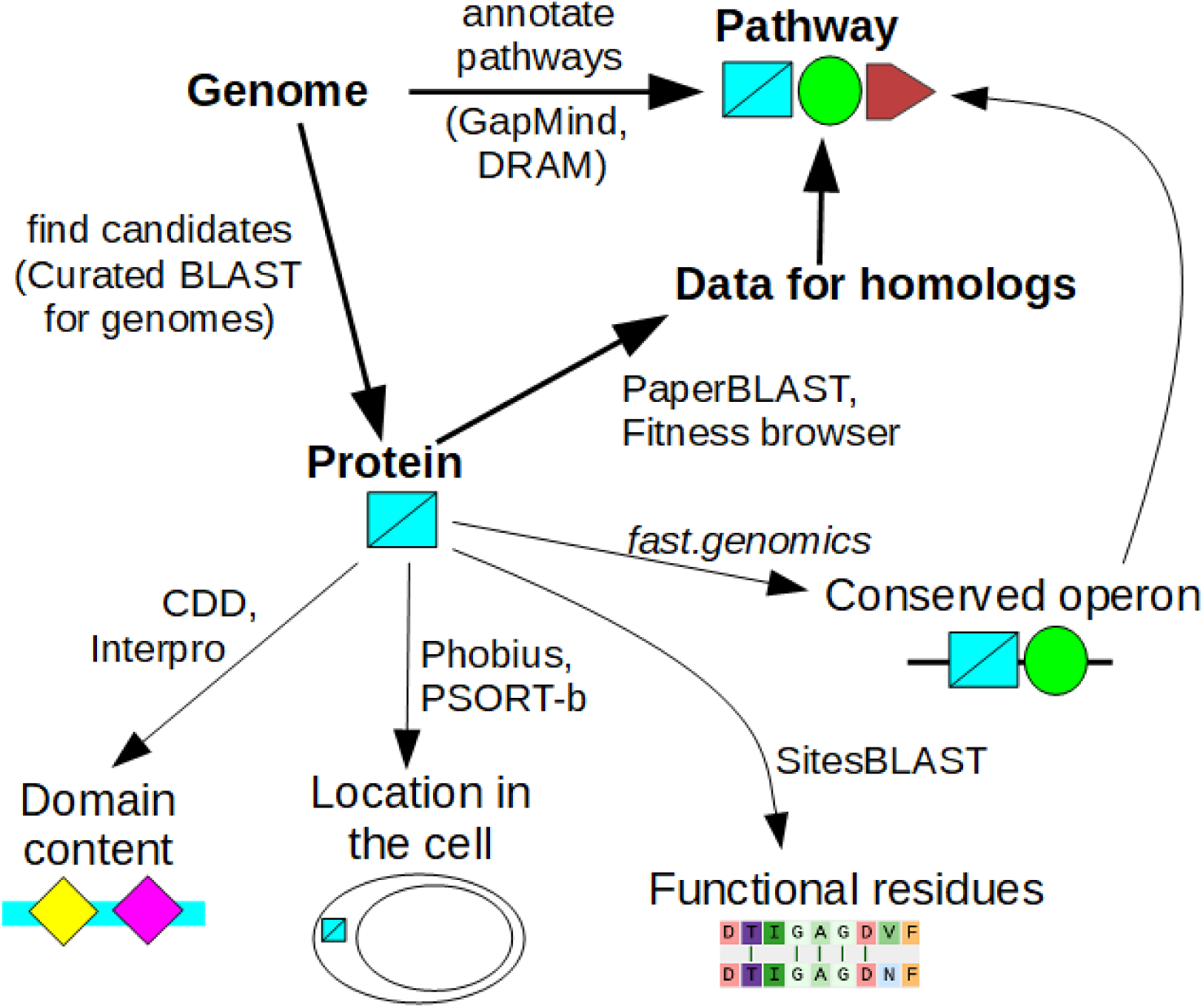
Overview of interactive tools. If you are interested in whether the genome encodes a specific capability, look at pathway annotations from GapMind or DRAM, or search for candidate proteins using Curated BLAST for genomes. Once you are interested in a specific protein, search for homologs of known function or for homologs with mutant phenotypes. If data for close homologs is available, this will often suggest a role for the protein. Otherwise, look for protein domains, predicted localization, known functional residues, and conserved operons. These approaches can give complementary hints as to the protein’s function.

This commentary will focus on enzymes and transporters, mostly because we have more experience with annotating metabolism than with other aspects of bacterial physiology. But these approaches apply to other types of proteins as well. In any case, accurate annotation of enzymes and transporters is central to understanding the metabolic capabilities of diverse bacteria.

## Results

### How accurate are gene annotations for enzymes and transporters?

To estimate the accuracy of gene annotations for enzymes and transporters, we needed a large set of metabolic genes of known function from diverse bacteria. We decided to focus on catabolic genes, as identified using randomly-barcoded transposon sequencing (RB-TnSeq, (Wetmore et al. 2015)). Catabolic genes can usually be identified from RB-TnSeq data because they should be important for fitness during growth with that specific carbon or nitrogen source, but not during growth in most other conditions. Furthermore, a specific molecular function can often be inferred from these mutant phenotypes (Price et al. 2018b). For example, if a transporter is specifically important during growth on L-fucose, it is probably an L-fucose transporter. Similarly, if a putative sugar kinase is specifically important during growth on D-glucosamine, then the kinase is probably either a glucosamine kinase or an N-acetylglucosamine kinase, depending on whether the organism catabolizes glucosamine via N-acetylglucosamine (Price et al. 2022); the correct pathway can usually be identified by examining the other genes that are specifically important for glucosamine utilization.

We selected a random sample of 500 genes with specific phenotypes on carbon or nitrogen sources from the Fitness Browser (http://fit.genomics.lbl.gov; (Price et al. 2018b)). A gene has a specific phenotype in an experiment if the abundance of its mutants changed by at least 2-fold, the change is statistically significant, and little change was seen in most other experiments (Price et al. 2018b). For each of these 500 genes, we manually determined if the phenotype appeared to be due to a role in the uptake or enzymatic breakdown of that carbon source or nitrogen source. We found 186 enzymes and transporters with specific roles in catabolism whose molecular function could be inferred using the genetic data. The other 314 genes either did not appear to encode enzymes or transporters, or the gene was detrimental to fitness, or the phenotype did not seem so specific, or we could not assign a specific molecular function in catabolism.

Of the proteins with inferred functions in catabolism, about half (96/186) had previously been reannotated in the Fitness Browser (Price et al. 2018b), but while the Fitness Browser’s reannotations are biased towards novel biology, the 186 proteins represent a random sample of the catabolic functions in the bacteria that we have RB-TnSeq data for. These proteins are from 31 different bacteria. Most of the proteins are from the phylum Pseudomonadota (α,β,γ-Proteobacteria), and about half (95/186) are from the genus *Pseudomonas*. These are well-studied taxa, so these proteins should be relatively easy to annotate.

We considered an automated annotation to be correct, or close enough, if it implied the same enzymatic or transport reaction to occur on the substrate that was implied by the mutant phenotype. If the mutant phenotypes implied multiple substrates, and the automated annotation included just one of those substrates, it was still deemed correct. Conversely, if the automated annotation listed additional substrates beyond what was expected from the mutant phenotypes, it was still considered correct: the protein may have other functions beyond those that were captured by the mutant phenotypes.

We considered the lack of a specific annotation to be an error, as a missing annotation could wrongly imply that the organism lacks that activity. In particular, vague annotations such as “sugar kinase” or “multiple sugar transport system” that did not mention any specific substrates were considered as errors, even if the substrate that was inferred from the mutant phenotypes was a sugar. If the tool did not report any annotation, that was also considered an error. Similarly, for fusion proteins with two entirely different activities, the annotation was considered incorrect if it only included one of those activities. (Three of the 186 catabolic proteins are fusion proteins.)

We tested four approaches for automated annotation. First, we used the annotation of the closest homolog from Swiss-Prot, which is a database of ∼570,000 curated annotations (UniProt Consortium 2023). Most of the Swiss-Prot annotations are based on homology, but a significant fraction are supported by experimental evidence. For Swiss-Prot homologs, we required the alignment to cover 70% of both proteins. Second, we used the ghostkoala tool from KEGG, which is based on another large database of curated annotations (Kanehisa et al. 2016a). Third, we used annotations from RefSeq, which are produced by NCBI’s Prokaryotic Genome Annotation Pipeline (PGAP, (Haft et al. 2024)). PGAP’s annotations are primarily based on curated descriptions of protein families, as defined by hidden markov models. Finally, we used CLEAN, which is a machine learning approach based on protein language models and contrastive learning (Yu et al. 2023). Since CLEAN is aimed at predicting enzyme function, we ran it on the 114 enzymes only.

If we consider vague or missing annotations to be errors, then annotation errors were common, with accuracies of 50-69% (Table 1). If we ignore vague or missing annotations, then the accuracy rises to 64-88%. Enzyme annotations were noticeably more accurate than transporter annotations (79% vs. 54% for Swiss-Prot best hits).

**Table 1:**
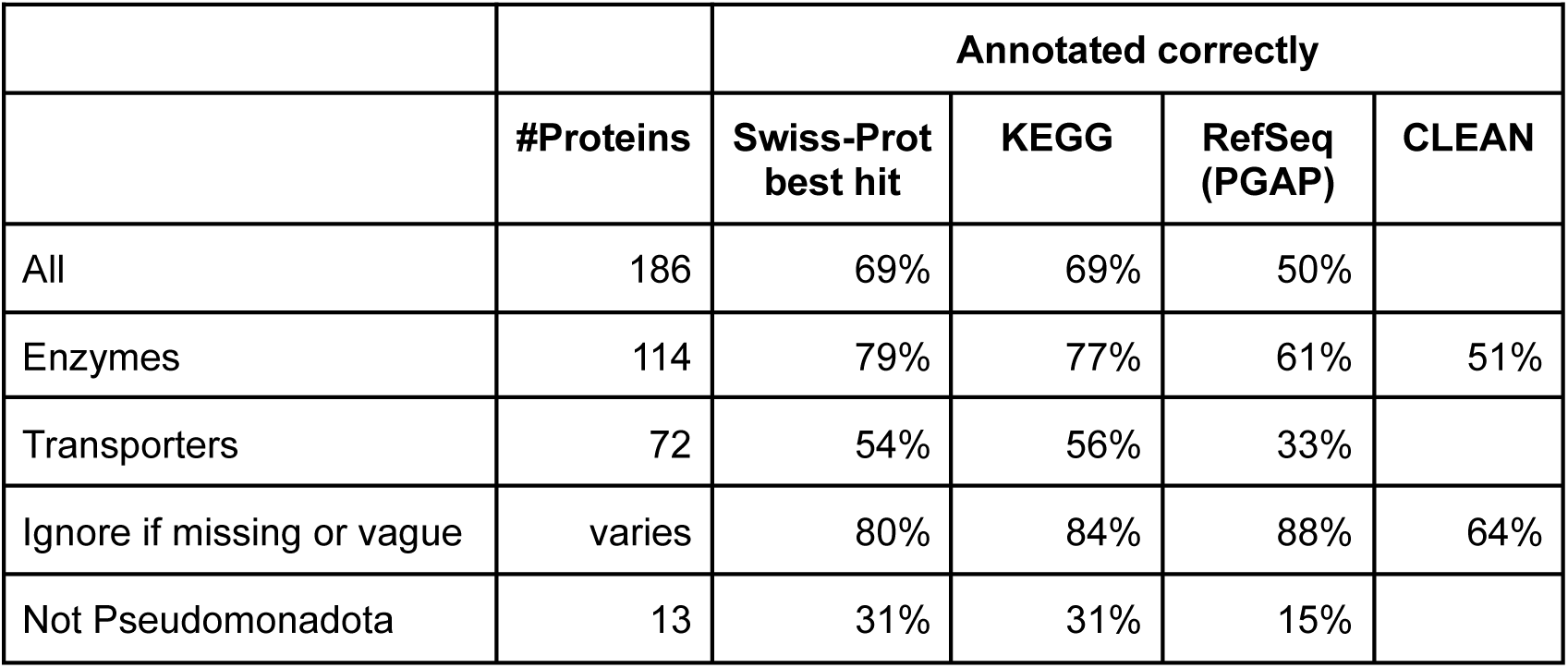
Accuracy of automated annotations for a random sample of catabolic enzymes and transporters whose function was inferred from mutant phenotypes.

|  |  | **Annotated correctly** |  |  |  |
| --- | --- | --- | --- | --- | --- |
|  | **#Proteins** | **Swiss-Prot best hit** | **KEGG** | **RefSeq (PGAP)** | **CLEAN** |
| All | 186 | 69% | 69% | 50% |  |
| Enzymes | 114 | 79% | 77% | 61% | 51% |
| Transporters | 72 | 54% | 56% | 33% |  |
| Ignore if missing or vague | varies | 80% | 84% | 88% | 64% |
| Not Pseudomonadota | 13 | 31% | 31% | 15% |  |

Not surprisingly, accuracy was higher when the annotation was transferred from a closer homolog. For Swiss-Prot, if the best homolog was 80% or identical or more, then the annotation was almost always accurate (46/47 proteins). Annotations from homologs that were 50-80% identical were mostly accurate (85%). But if the best homolog in Swiss-Prot had under 50% identity, transferring the annotation from the best hit was accurate just 61% of the time. Similarly, enzyme annotations from CLEAN were far more accurate if they were labeled as high-confidence (91% vs. 34% otherwise).

The protein with a different function than expected given its close homolog in Swiss-Prot was HSERO_RS05250, which is a transporter subunit that is specifically important for L-fucose utilization (Figure 2A). HSERO_RS05250 is 80% identical to A0B297, which Swiss-Prot annotates as acting on ribose, galactose, or methyl galactoside. Using PaperBLAST (Price and Arkin 2017), we could not find any experimental data about the function of close homologs of HSERO_RS05250 or A0B297 besides the RB-TnSeq data. This subfamily is usually encoded near fucose catabolism genes such as L-fucose dehydrogenase, L-fucono-1,5-lactonase, and L-fuconate dehydratase (Figure 2B). So we believe that A0B297’s annotation in Swiss-Prot is erroneous. The misannotation of A0B297 is probably based on the experimentally-supported functions of the RbsA and MglA proteins of *Escherichia coli*, which are 44-47% identical to A0B297.

**Figure 2:**
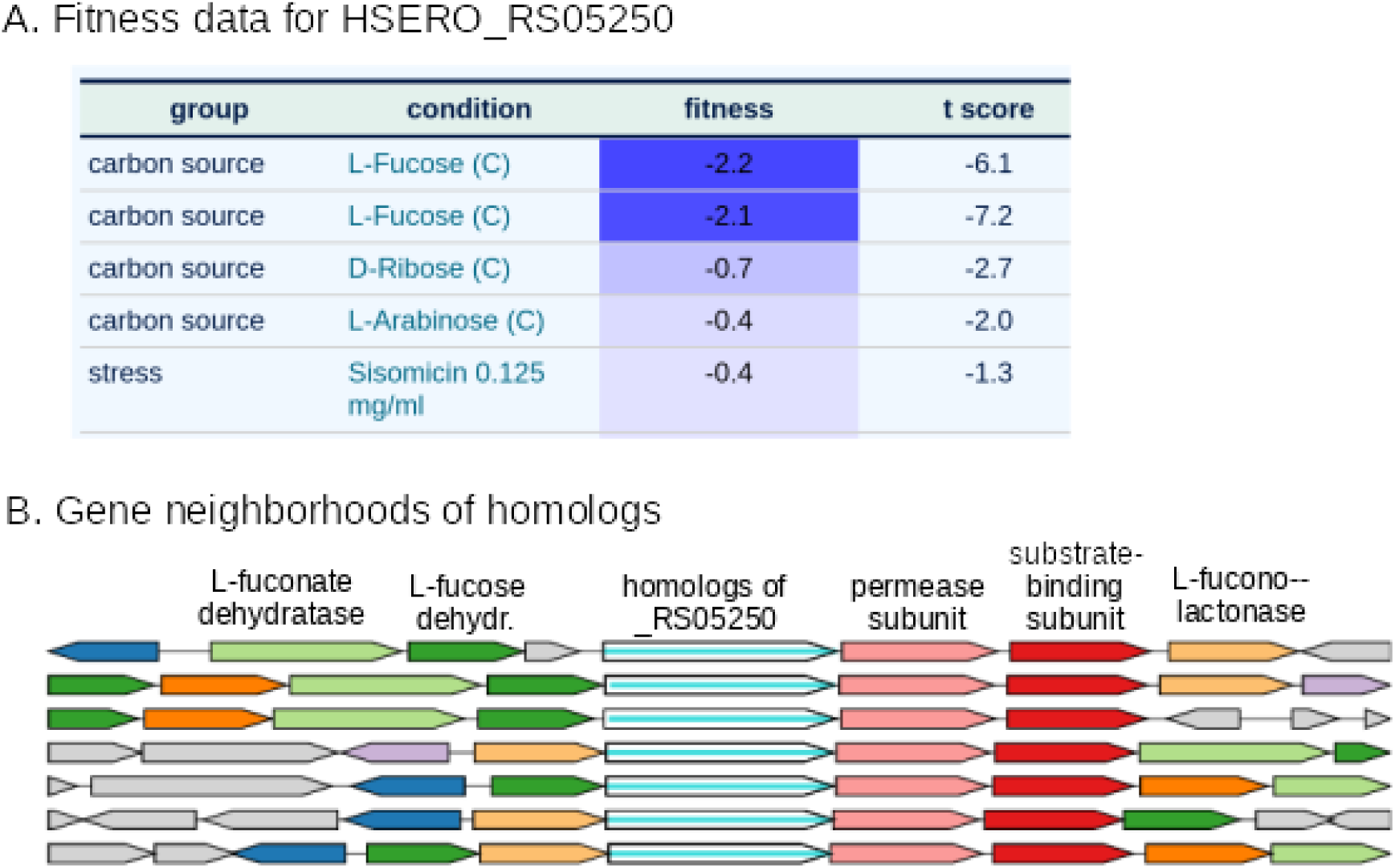
Identification of a L-fucose transporter using RB-TnSeq data and conserved gene neighbors. (A) HSERO_RS05250 is specifically important for L-fucose utilization. This screenshot from the Fitness Browser shows the strongest fitness values for this gene, from 96 experiments. A gene’s fitness is the log2 change in the relative abundance of its mutants during an experiment (such as growth in L-fucose). Negative fitness values indicate that mutants are at a disadvantage. (B) Gene neighbors of homologs of HSERO_RS05250 confirm that it is involved in fucose catabolism. This screenshot from *fast.genomics* shows the gene neighborhoods for seven homologous proteins (72-100% identity) from different genera. Genes are color-coded if similar proteins are present in more than one track. The labels at top with the likely function of each gene were added by hand. L-fucose dehydrogenase, L-fucono-1,5-lactonase, and L-fuconate dehydratase are the initial steps in an oxidative pathway for fucose catabolism (Hobbs et al. 2013). HSERO_RS05250 is the ATPase subunit of an ABC transporter. The putative permease and substrate-binding subunits are also encoded nearby.

The accuracy of the automated annotations dropped dramatically if we excluded proteins from Pseudomonadota, which is the best-studied phylum of bacteria. The remaining proteins are from Bacteroidota (10 proteins) or Desulfobacterota (3 proteins). Annotation accuracy for these other proteins was just 31%, either for Swiss-Prot best hits or for KEGG. The poor accuracy reflects the lower level of study of those phyla, which leads to a lower similarity of their proteins to characterized proteins. For proteins with homologs in Swiss-Prot, proteins from Pseudomonadota had a median identity to their best hit of 60%, while proteins from other phyla had a median identity of just 39%.

### How up-to-date are automated gene annotations?

The low accuracy of annotation of these 186 proteins is ironic given that we had previously published the functions for about half of them, and that most of the relevant RB-TnSeq data was published over five years ago (Price et al. 2018b). However, as of December 2023, none of these 186 proteins’ functions have been updated based on the RB-TnSeq data, either in Swiss-Prot or in other curated resources of metabolism, such as MetaCyc (Caspi et al. 2020) or BRENDA (Chang et al. 2021).

The low rate of curation of genes whose function was inferred using RB-TnSeq could reflect skepticism about our approach. To get a broader estimate of the rate at which papers about protein functions are curated, we considered a sample of 32 papers, published in mBio in late 2016, that make claims about protein function in their abstract. (This is the subset of the 54 papers analyzed in (Price and Arkin 2017) that mention the protein’s function in the abstract.) As of January 2024, none of these 32 papers are referred to by curated Swiss-Prot entries. We then asked whether the key proteins were listed as characterized in Swiss-Prot or in other curated databases of experimentally-characterized proteins that are incorporated into PaperBLAST (Table 2). We considered not only the original paper, but also whether other papers about those proteins’ function had been curated. We found that in 16 cases, the protein(s), or nearly-identical orthologs from another strain of the same species, were curated. Overall, in over seven years, half of the characterized proteins have been curated.

**Table 2:** Sources of experimentally-characterized proteins in the PaperBLAST database.

| Database | Topic | Characterized proteins | Reference |
| --- | --- | --- | --- |
| Swiss-Prot | all proteins | 102,037 | (UniProt Consortium 2023) |
| BRENDA | enzymes | 29,304 | (Chang et al. 2021) |
| MetaCyc | metabolism | 12,664 | (Caspi et al. 2020) |
| CAZy | carbohydrate metabolism | 8,629 | (Lombard et al. 2014) |
| TCDB | transporters | 8,505 | (Saier et al. 2016) |
| CharProtDB | characterized proteins | 7,173 | (Madupu et al. 2012) |
| REBASE | restriction enzymes | 3,818 | (Roberts et al. 2015) |
| EcoCyc | *E. coli* | 3,261 | (Karp et al. 2023) |
| Fitness Browser reannotations | mutant phenotypes | 1,757 | (Price et al. 2018b) |
| PRODORIC | transcription factors | 345 | (Dudek and Jahn 2022) |

The low rate of curation reflects the limited resources available. Swiss-Prot has roughly three full-time biocurators for all prokaryotic proteins, and “tens of thousands of publications remain to be curated for enzymes” (de Crécy-Lagard et al. 2022).

Automated annotations may be even more out of date, relative to current knowledge, because the underlying reference databases may be years out of date. Two of the most popular tools for annotating bacterial genomes are Prokka and the RAST server: each has over 10,000 citations in Google Scholar (Aziz et al. 2008; Seemann 2014). As of January 2024, Prokka’s reference database is based on the October 2019 release of Swiss-Prot. The RAST server’s reference databases date to 2016 or earlier (Davis et al. 2016).

Finally, in practice, protein annotations are often obtained from RefSeq, Genbank, or TrEMBL (the non-curated part of UniProt). RefSeq updates their annotation pipeline (PGAP) and recomputes annotations every so often. but annotations in Genbank or TReMBL are rarely updated. Furthermore, annotations in Genbank or TReMBL have high error rates, probably much higher than for the automated approaches that we considered (Schnoes et al. 2009; Rembeza and Engqvist 2021).

### Why is automated annotation difficult?

Most enzymes and transporters probably belong to known families. For instance, of the 186 catabolic proteins discussed above, 172 (92%) have homologs in Swiss-Prot, and even the vague annotations indicate a type of reaction (i.e., “uncharacterized oxidoreductase” or “sugar kinase”) or that the protein is a transporter. Just 10 of the 186 proteins (5%) could not be annotated as the correct type of enzyme, or as a transporter, by using the best hit from Swiss-Prot. However, if a protein from a known family is not closely related to any characterized protein, it is difficult to identify the correct substrate.

To quantify what fraction of bacterial proteins are similar to characterized proteins, we began with a random sample of 2,000 protein-coding genes from diverse bacteria. These were taken from representative genomes in *fast.genomics (Price and Arkin 2024)*. We compared these proteins to a database of 189,323 experimentally-characterized proteins from all kingdoms of life, which was compiled from ten different databases in the December 2023 release of PaperBLAST (Table 2). We used ublast (Edgar 2010) to find alignments with an expectation value of 10^-10^ or less and required that the alignment cover 70% of both proteins. We found that just 28% of bacterial proteins are ≥40% identical to a characterized protein. Even at around 40% identity, annotations for enzymes or transporters will have significant error rates. For instance, for catabolic proteins that were 35%-50% identical to their best hit in Swiss-Prot, and for which the best hit had a specific annotation, the error rate was 34%.

Of the bacterial proteins in our sample, most are from genomes of isolates, but 19% are from high-quality metagenome-assembled genomes (MAGs). (*Fast.genomics* uses a MAG as the representative for a genus if it is high quality and if no high-quality isolate genome is available.) Not surprisingly, proteins from bacterial MAGs are less likely to be ≥40% identical to a characterized protein than proteins from bacterial isolates are (21% vs. 30%, P = 0.0004, Fisher exact test).

Archaeal proteins are less likely than bacterial proteins to be similar to a characterized protein. We compared 2,000 random protein-coding genes from diverse archaea (again from representative genomes in *fast.genomics*) to our database of characterized proteins. Just 21% of archaeal proteins are ≥40% identical to a characterized protein.

## Discussion

Overall, most prokaryotic proteins do not have a close characterized homolog in the curated databases, which makes it difficult to predict their functions accurately. Hence the need for interactive tools.

### Finding characterized homologs with PaperBLAST

To determine the likely function of a protein, our first stop is always PaperBLAST, which finds papers about homologs (http://papers.genomics.lbl.gov/; (Price and Arkin 2017)). PaperBLAST takes just a few seconds to compare the query to proteins from 14 curated databases as well as to proteins that are mentioned in scientific articles (Table 3). Ideally, a PaperBLAST search will lead to experimental data about a homolog with a known function, either from one of the curated databases, or from a paper that has not been curated yet. As of a few years ago, the odds of finding useful information in PaperBLAST about a vaguely-annotated bacterial protein was around 22% (Price and Arkin 2017). However, we recommend using PaperBLAST for all proteins, as it can quickly reveal obvious errors in more-specific annotations, such as the L-fucose transporter discussed above.

**Table 3:**
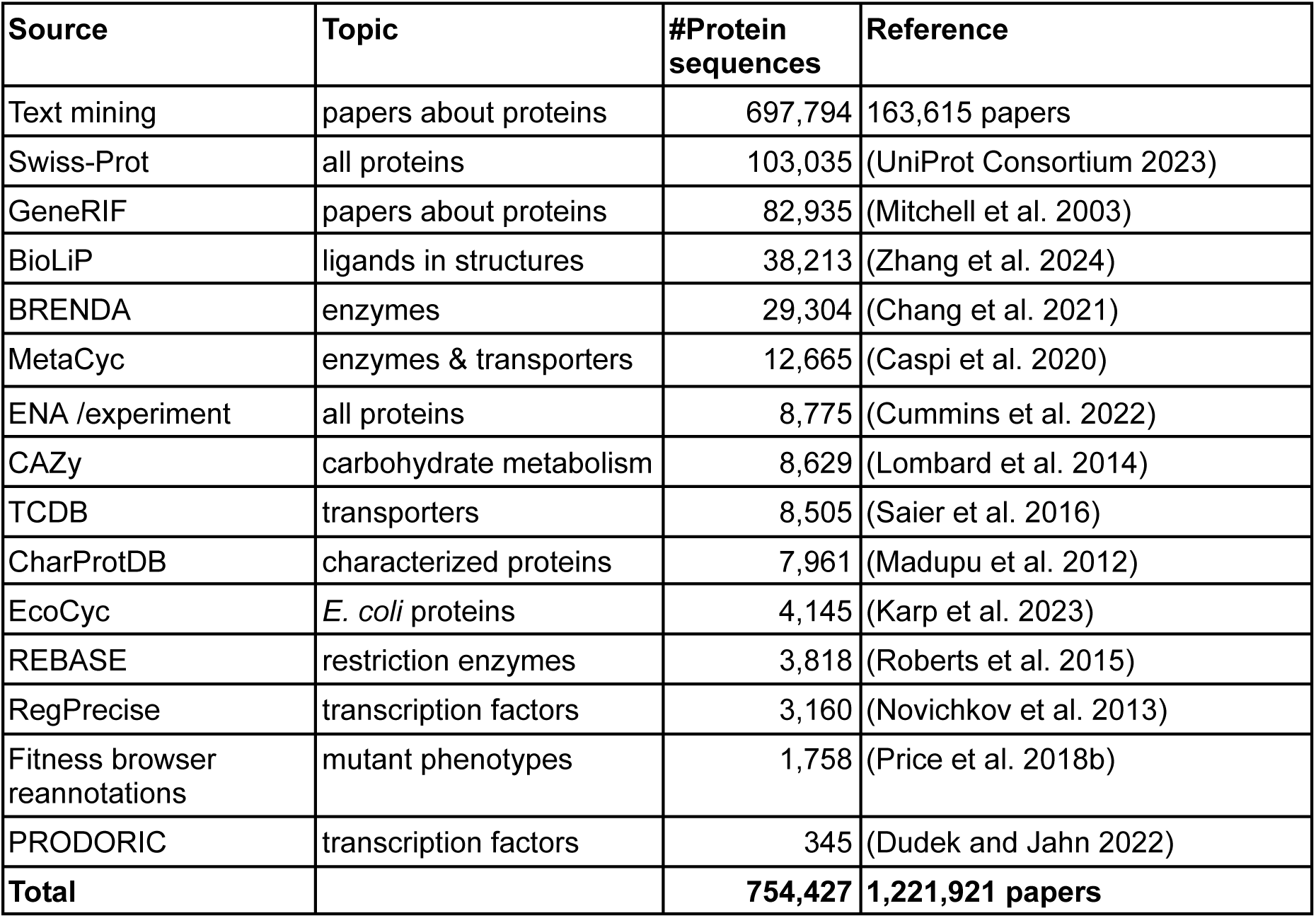
Data sources for the PaperBLAST database. For databases that include experimentally-characterized proteins, counts may be higher than in Table 2 because a few proteins have functionally-uninformative annotations, such as “DUF3828 domain-containing protein YqhG”. Except for EcoCyc and RegPrecise, only proteins with experimental evidence or literature references are included. For example, 18% of Swiss-Prot entries have experimental evidence and are included. For the European Nucleotide Archive (ENA), annotations with the “experiment” tag are further filtered, see Materials and Methods. PaperBLAST’s text mining uses EuropePMC (Europe PMC Consortium 2015) to search scientific articles, including the full text of most articles, for protein identifiers.

As another example, consider the dehydrogenase Shewana3_2071, which was one of the catabolic proteins in our annotation test. The best hit in Swiss-Prot is a sulfoquinovose 1-dehydrogenase (P0DOV5, 46% identity). but Shewana3_2071 is specifically important for L-arabinose utilization. As shown in Figure 3, several of the top results from PaperBLAST are informative as to its function. The top result is the reannotation from the Fitness Browser, based on the mutant phenotype. The second result is the crystal structure of a similar protein in the complex with NADH; this confirms that this is a family of dehydrogenases, but does not have direct information as to the other substrate. The third result is a paper about C785_RS21245 from *Herbaspirillum huttiense*; the snippet in PaperBLAST’s results mentions functional characterization. That paper shows that the homolog is a 1-aldose dehydrogenase, with a higher specificity constant (k_cat_/K_M_) for L-arabinose than for other plausible substrates (Watanabe et al. 2019). (The specificity constant for D-fucose was 3-fold higher than for L-arabinose, but D-fucose is a rare compound in nature.) The similarity of Shewana3_2071 (54% identity) to a biochemically-characterized L-arabinose dehydrogenase (which was published after we made this reannotation) confirms that Shewana3_2071 is an L-arabinose dehydrogenase. On the other hand, the next-best substrate of C785_RS21245, D-xylose, has a specificity constant that is only 14-fold lower than for L-arabinose. C785_RS21245 itself probably does not contribute significantly to D-xylose degradation: *H. huttiense* has a different D-xylose-specific dehydrogenase whose expression is induced by D-xylose (Watanabe et al. 2019). But in a different genetic context, C785_RS21245 probably could function in D-xylose degradation. This illustrates how enzymes’ annotations often simplify their biochemical capabilities.

**Figure 3:**
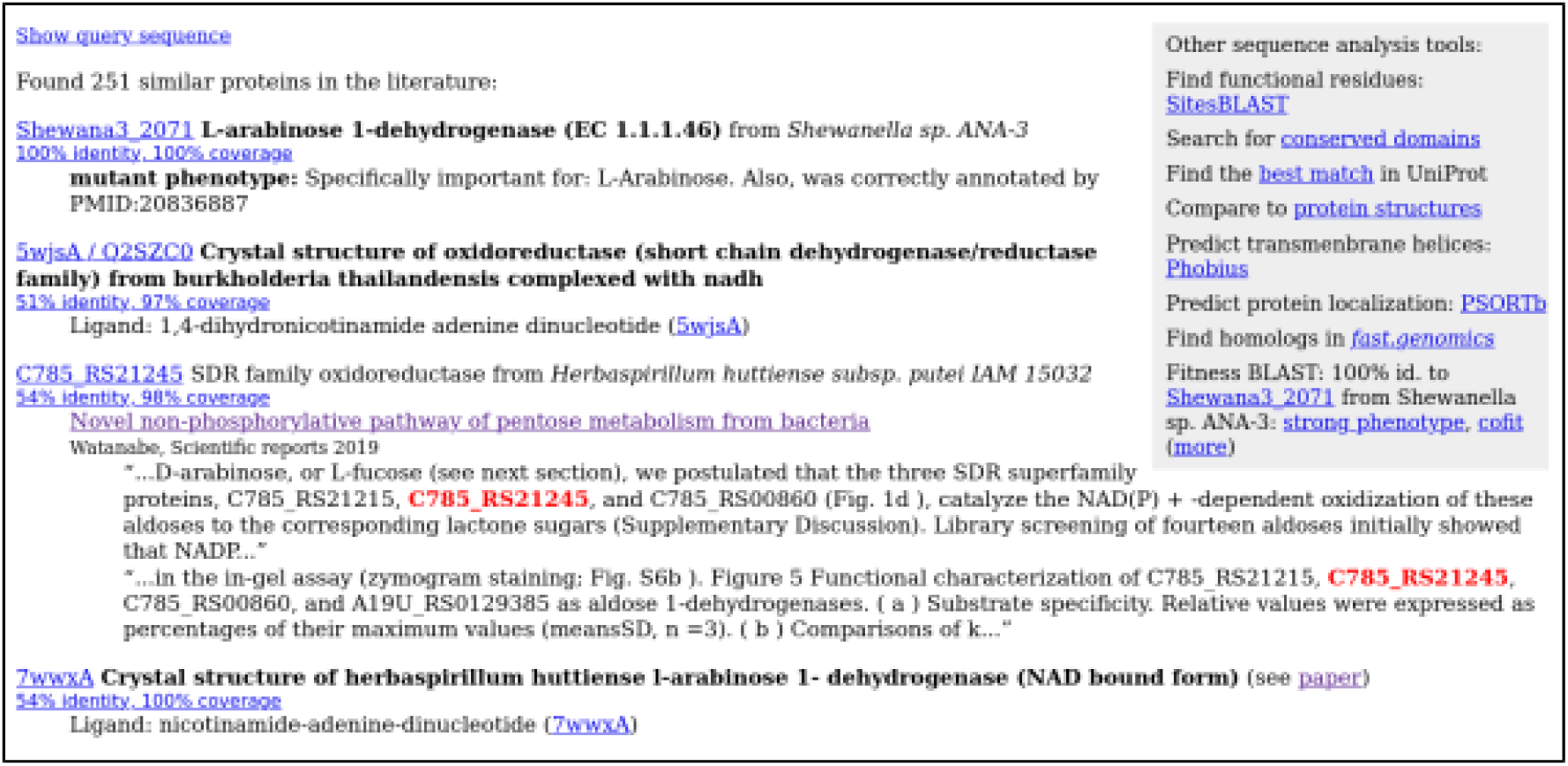
Finding papers about a protein and its homologs with PaperBLAST. This screenshot shows the top results for Shewana3_2071. Experimentally-characterized homologs are shown with curated annotations in bold. Homologs that are discussed in papers are shown with snippets of text from those papers. Within each snippet, the homolog’s identifier is highlighted. The top right shows links to other interactive tools.

To infer a protein’s function from a characterized homolog, the alignment should cover almost the full length of both sequences. If not, examine the domain content (see below). If the proteins are similar over their full length, or contain the same domains, then how close is close enough to infer that the protein of interest has the same function as the characterized homolog? As a rule of thumb, for enzymes, the most similar characterized sequence is likely to have the same function if it is above 40% identity. (For example, in the annotation test, for enzymes whose best hit in Swiss-Prot was 35-50% identical and had a specific annotation, the annotation was correct 72% of the time.) It is likely to have a similar function (but perhaps with a somewhat different substrate) if above 30% identity. As discussed above, transporters are more difficult to annotate, and their specificity evolves more quickly, so a higher %identity is needed to reach the same confidence.

We update the PaperBLAST database every two months, so it is much more up-to-date than is possible with automated annotations. Specifically, for each update, we rerun PaperBLAST’s text mining, and we update its copies of Swiss-Prot, GeneRIF, BioLiP, EcoCyc, and the Fitness Browser reannotations. We don’t update the other curated databases in PaperBLAST as frequently. Also, PaperBLAST’s copy of MetaCyc is not being updated because it is no longer freely available, and CharProtDB is no longer being updated.

### Viewing mutant phenotypes in the Fitness Browser

PaperBLAST also shows any similarity to proteins with mutant phenotypes from the Fitness Browser. We call this “Fitness BLAST” (see far right of Figure 3). In particular, PaperBLAST highlights if a homolog has a strong phenotype in a condition (fitness under −2, meaning that mutants decreased in abundance by 4-fold or more), or if it has a similar fitness pattern as other genes (“cofitness”).

The Fitness Browser includes randomly-barcoded transposon data (RB-TnSeq) from diverse bacteria and archaea (Price et al. 2018b). As of February 2024, the Fitness Browser contains 7,552 genome-wide experiments from 46 bacteria and two archaea. Most of these bacteria are from the phylum Pseudomonadota, but the Fitness Browser also includes data for four Bacteroidota, two Desulfurbacterota, one Actinomycetota, and one Cyanobacteriota.

We consider a protein to have a functional link, based on its mutant phenotypes, if it has a specific phenotype (as defined above, see Figure 2A for an example) or if it is sufficiently cofit with another gene. Specific phenotypes or cofitness often indicate a functional relationship, especially if they are conserved across similar proteins (Price et al. 2018b). To quantify the cofitness of two genes, we use the linear correlation between their fitness profiles. Sufficient cofitness was defined as a correlation of at least 0.8, or a correlation of at least 0.6 and similar proteins in another bacterium have cofitness at least 0.6 (Price et al. 2018b). Overall, the Fitness Browser contains functional links for 33,261 bacterial proteins. Across the random sample of 2,000 proteins from diverse bacteria, 25% have a homolog with a functional link that is over 40% identity, with 70% coverage both ways.

To go from a specific phenotype for a putative enzyme to the correct substrate, it is often necessary to consider the rest of the pathway. For example, consider the putative sugar kinase SM_b21217 from *Sinorhizobium meliloti*, which is specifically important for growth with glucosamine as the nitrogen source. Depending on which pathway *S. meliloti* uses to consume glucosamine, SM_b21217 might be a glucosamine kinase or an N-acetylglucosamine kinase (Figure 4A; (Price et al. 2022)). If *S. meliloti* uses the acetylated pathway, then a transacetylase and a deacetylase would also be involved in glucosamine utilization, and the same transporter and the same kinase might be used during utilization of both glucosamine and N-acetylglucosamine (Figure 4A). The relevant proteins can be found by homology (see Curated BLAST for genomes below), or by using the fitness data. In particular, by examining the genes that are important for utilization of N-acetylglucosamine but not glucosamine as the nitrogen source, we can see that N-acetylglucosamine kinase and a transporter are important for the utilization of N-acetylglucosamine only (Figure 4B). This suggests that *S. meliloti* does not use the acetylated pathway for glucosamine utilization, and hence that SM_b21217 is a glucosamine kinase. (Another test of which pathway *S. meliloti* uses would be to search for the glucosamine N-acetyltransferase using Curated BLAST for genomes, which is described below.)

**Figure 4:**
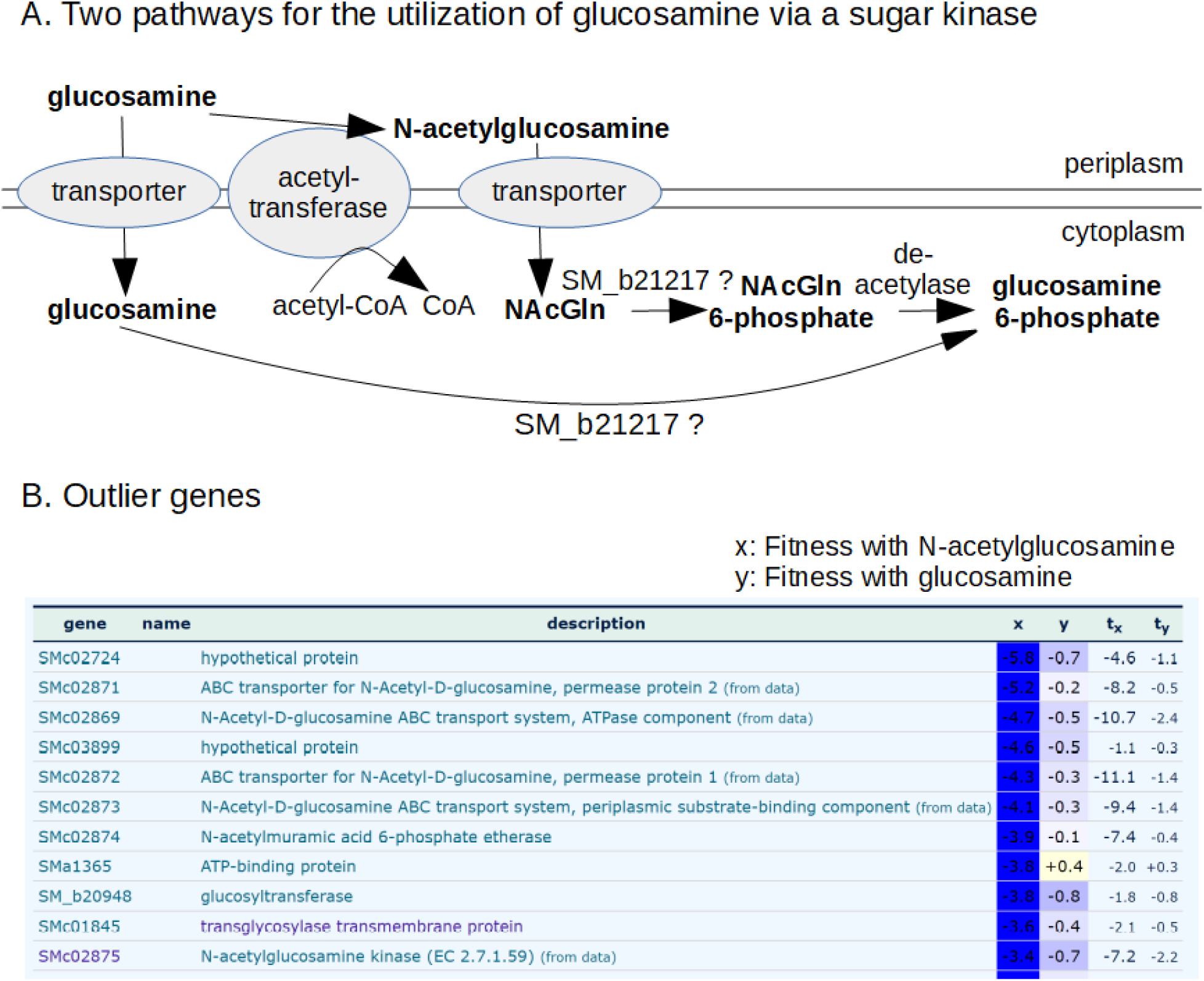
Using the Fitness Browser to consider alternate pathways. (A) Two potential roles for the putative sugar kinase SM_b21217 in glucosamine utilization (adapted from (Price et al. 2022)). (B) A comparison of fitness data from *Sinorhizobium meliloti* with N-acetyglucosamine or glucosamine as the nitrogen source. In this screenshot from the Fitness Browser, genes that are important for utilization of N-acetyglucosamine, but not glucosamine, are listed.

More broadly, reasoning about the expected phenotypes for a protein will often require examining which other functionally-related proteins are present and what their phenotypes are. To support this style of reasoning, the Fitness Browser can render the fitness data in many different ways, including scatterplots for comparing genes’ fitness values, lists of cofit genes, lists of genes with specific phenotypes in a condition, scatterplots for comparing experiments, lists of outlier genes (as in Figure 4B), heatmaps, and comparisons of the fitness data for similar proteins from different bacteria in the same condition. A related issue that often arises is isozymes. If there are two isozymes for one step in the pathway, then they may be genetically redundant, so that neither gene shows the expected phenotype.

One caveat with RB-TnSeq data is the possibility of polar effects: transposon insertions within a gene that is in an operon may disrupt the expression of a downstream gene. If this occurs, then insertions in the gene can have a strong phenotype even though only the downstream gene is important for fitness in that condition. Polar effects are not predominant in the RB-TnSeq data, but they are not rare either (Wetmore et al. 2015). A common sign that a phenotype is due to a polar effect is that the effect depends on which orientation the transposon inserted in. If the antibiotic resistance marker’s promoter is in the same orientation as the disrupted gene, then the promoter can drive expression of the downstream genes, so polar effects usually occur only for insertions in the opposite orientation. In most pages in the Fitness Browser, clicking on a gene fitness value will show the strain fitness values for insertions in and around the gene, along with the orientation of the antibiotic resistance marker in each strain. Because strain fitness values are much noisier than gene fitness values, the Fitness Browser makes it easy to average the strain fitness value across replicate experiments.

Finally, in our experience, inferring a genes’ likely function from its phenotypes is more straightforward for metabolic genes and transporters than for other types of proteins. Many proteins have pleiotropic phenotypes that are difficult to rationalize, even if the protein is well-characterized.

### Checking a protein’s domain content or finding distant homologs

If homologs of known function are not available, or if the alignments do not cover the whole sequence (which suggests that the domain structure may have changed), then we recommend examining the domain structure of the protein of interest, using the Conserved Domain Database (CDD, (Marchler-Bauer et al. 2015)) or InterPro (Finn et al. 2017). Searching CDD is fast and can be run interactively. Analyzing a sequence with InterPro is much slower, but pre-computed results for all of UniProt (over 250 million proteins) are available. For proteins from public bacterial genomes, an identical or nearly-identical homolog is probably in UniProt, and these can be found quickly using SANSparallel (Somervuo and Holm 2015). Links to CDD and to UniProt searches are included in the PaperBLAST results. Either CDD or InterPro will show any similarity of the protein sequence to profiles (hidden markov models) of protein families, such as from PFam (Finn et al. 2014). Profile comparisons are more sensitive than pairwise sequence comparisons, so these tools often find homology that are missed by BLAST-based tools.

Any domains that are present in the characterized homolog but missing from the protein of interest might indicate a loss of function. For example, the characterized protein might have two enzymatic activities and the protein of interest might have kept just one of them. Conversely, any additional domains in the protein of interest could indicate an additional function. Although domain content is very useful for generating hypotheses about a protein’s function, many domains or families are so broad that they include proteins with very different functions.

Another approach to finding distant homologs is to use the predicted structure and to search for proteins of similar structure. AlphaFold predictions are available for most of the proteins in UniProt (Varadi et al. 2022), and FoldSeek can quickly compare these predictions to other structures and find remote homologs (van Kempen et al. 2024). Alternatively, if reliable BLAST-level homologs (down to 30% identity or a bit less) are available, then the Jackhmmer tool from the HMMer web server can build a profile from the closer homologs and then use that profile to find more distant homologs (Potter et al. 2018).

If a remote homolog with a known function is found using a tool like FoldSeek, it is interesting to see if the functional residues are conserved. For help finding functional residues, see SitesBLAST below. For aligning distant homologs, we recommend RCSB’s pairwise structure alignment tool (https://www.rcsb.org/alignment; (Burley et al. 2021)).

### Predicting a protein’s location in the cell

Another important aspect of a protein’s function is its location in the cell. In particular, the break-down of oligosaccharides often begins outside of the cell or in the periplasm. Smaller sugars may also be oxidized or cleaved in the periplasm. Conversely, most other catabolic reactions, and most biosynthetic reactions, take place in the cytoplasm. Understanding where the metabolic enzymes are located is also critical for correctly understanding the role of transporters (i.e., Figure 4A).

To predict a protein’s localization, we recommend using Phobius (Käll et al. 2004), which predicts signal peptides and trans-membrane helices, and PSORTb, which uses a variety of factors including the localization of homologs to predict a protein’s localization (Yu et al. 2010). Links to both tools are included in the PaperBLAST results page.

### Finding functional residues with SitesBLAST

Although highly-similar sequences are likely to have the same function, an enzyme’s function is not determined by %identity. Rather, most of the protein’s sequence serves to fold the protein into the correct overall shape, and the function is determined by a small number of residues that bind the substrate or participate in catalysis (sometimes called “active site” residues). Unfortunately, most of the popular automated tools for protein annotation do not consider functional residues. (The only exception we are aware of is UniProt’s UniRule (MacDougall et al. 2021).)

Information about these functional residues may be available from SitesBLAST, which makes it easy to see if the functional residues are conserved (Price and Arkin 2022). For example, the SitesBLAST results for the glucosamine kinase SM_b21217 shows that all of the active site residues, and most of the ADP-binding residues, are conserved (Figure 5). This confirms that SM_b21217 is a kinase.

**Figure 5:**
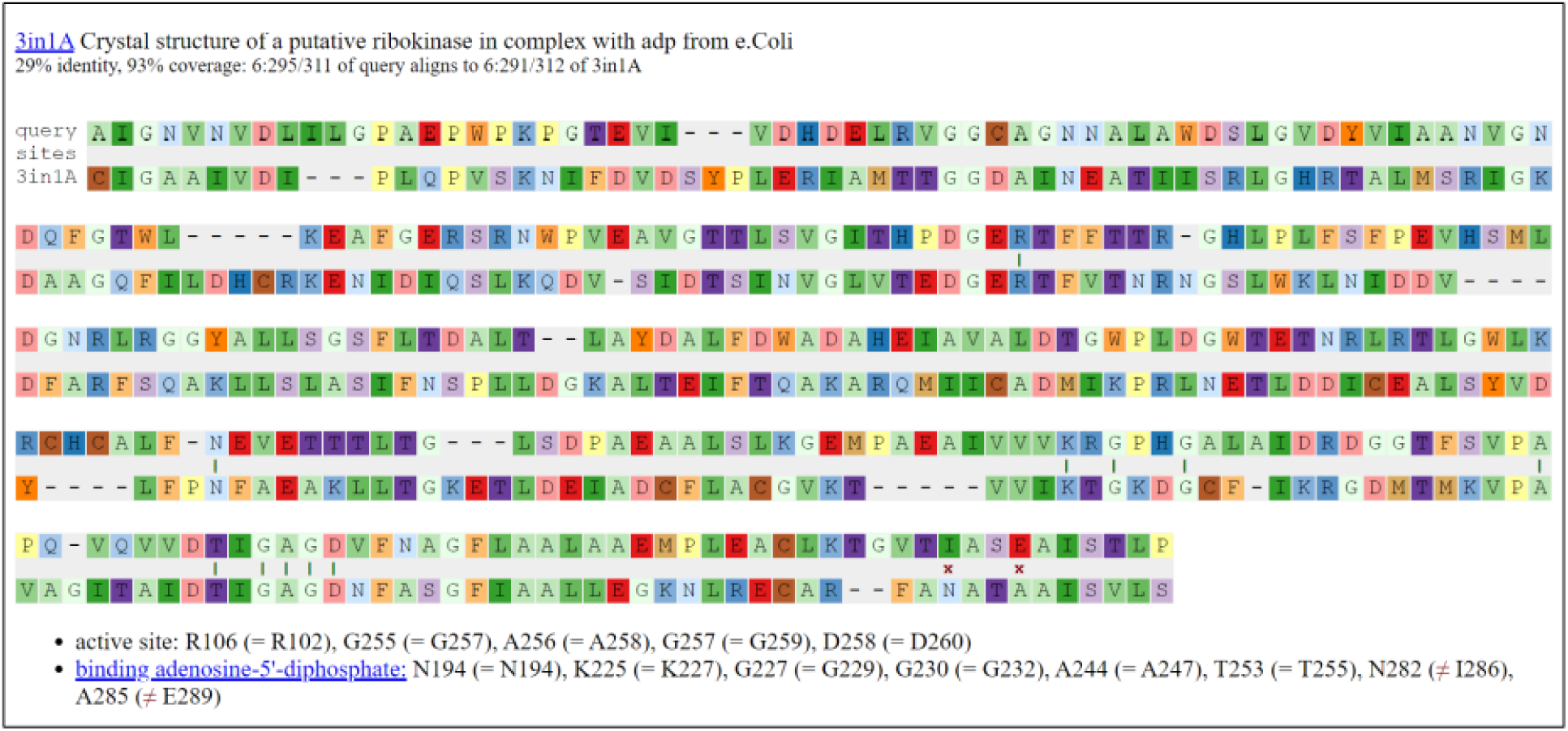
Checking functional residues with SitesBLAST. This screenshot shows the top result for the putative sugar kinase SM_b21217.

SitesBLAST takes just a few seconds and compares the input sequence to a database of 170,045 proteins with known functional residues. These functional residues include ligand-binding and active-site residues in crystal structures, as compiled by the BioLiP database (Zhang et al. 2024), and Swiss-Prot features with experimental evidence (UniProt Consortium 2023). SitesBLAST can identify potential functional residues for around half of all proteins (Price and Arkin 2022). SitesBLAST is available from the PaperBLAST results page or at http://papers.genomics.lbl.gov/sites.

If SitesBLAST shows that the catalytic residues are not exactly conserved, but you still suspect that the protein might be catalytically active, then we recommend checking the Mechanism and Catalytic Site Atlas (M-CSA; (Ribeiro et al. 2020)). This resource describes the catalytic mechanisms of hundreds of different enzymes, which can help you reason about the potential activity of a protein.

SitesBLAST compares just two proteins. If you want to compare functional residues across many proteins, then we recommend a related tool, Sites on a Tree (Price and Arkin 2022). Sites on a Tree can indicate whether changes to functional residues are compatible with a conserved function, or can identify subfamilies that are likely to have different functions.

### Testing a protein’s function with its structure

If you have a specific hypothesis about the protein’s function, you may be able to use the predicted structure to test your hypothesis. Structural analysis methods are computationally intensive and so are usually not interactive.

Predicted protein complexes can be checked by running AlphaFold on two or three proteins together. We use Google Colab Pro+ to run ColabFold (Mirdita et al. 2022). (No programming is required.) If there are high-confidence contacts between two protein chains, then they probably do form a complex (Yin et al. 2022).

Docking methods such as AutoDock Vina (Trott and Olson 2010) can in principle be used to predict whether a small molecule binds and where, and hence to constrain the substrate specificity of an enzyme or transporter. We are not aware of any large-scale tests of this approach for predicting specificity, and we have not had success with it ourselves. But identifying inhibitors or ligands by docking to AlphaFold models is not very accurate (Wong et al. 2022; Lyu et al. 2023). Even predicting how a given small molecule will bind a protein remains challenging, with the correct binding pose identified less than half the time (Krishna et al. 2023). So it appears that it is not (yet?) possible to predict small-molecule substrates accurately.

### Inferring function from gene neighborhoods or gene presence/absence

Another way to generate hints about a protein’s function is to examine the genes encoded near homologs of the protein of interest. In particular, genes in conserved operons often have related functions (Dandekar et al. 1998; Huynen et al. 2000; Wolf et al. 2001). The gene neighborhoods of homologs of a protein of interest can be viewed using *fast.genomics* (http://fast.genomics.lbl.gov/; (Price and Arkin 2024)). A conserved operon will show up as a group of genes that are encoded on the same strand and with a close spacing (usually under 100 nt) across multiple genera (see example in Figure 2B).

If you have a specific hypothesis about a protein’s function, then which genomes it is found in may also be informative. For instance, an unusual type of methionine synthase was reported in *Methanothermobacter marburgensis* (Schröder and Thauer 1999). We will call it MesA (Price et al. 2021). *In vitro*, MesA uses methylcobalamin instead of methyl-tetrahydrofolate as its methyl donor, but with a high K_M_ (Michealis-Menten constant); it is expected that a corrinoid (cobalamin-binding) protein would be the physiological donor (Schröder and Thauer 1999). We noticed that MesA is found only in methanogens (Price et al. 2021), which suggests that MtrA (the corrinoid subunit of tetrahydromethanopterin S-methyltransferase) might be the donor protein. As shown in Figure 6, genomes that contain reasonably close homologs of MesA, with a bit score above 30% of maximum, almost always contain MtrA. Also, in a few genomes, the two genes are encoded near each other and on the same strand (green points in Figure 6). This supports a functional relationship between MtrA and MesA. Some genomes with MtrA (score ratio above 0.4) lack MesA; these encode another type of methionine synthase (Price et al. 2021).

**Figure 6:**
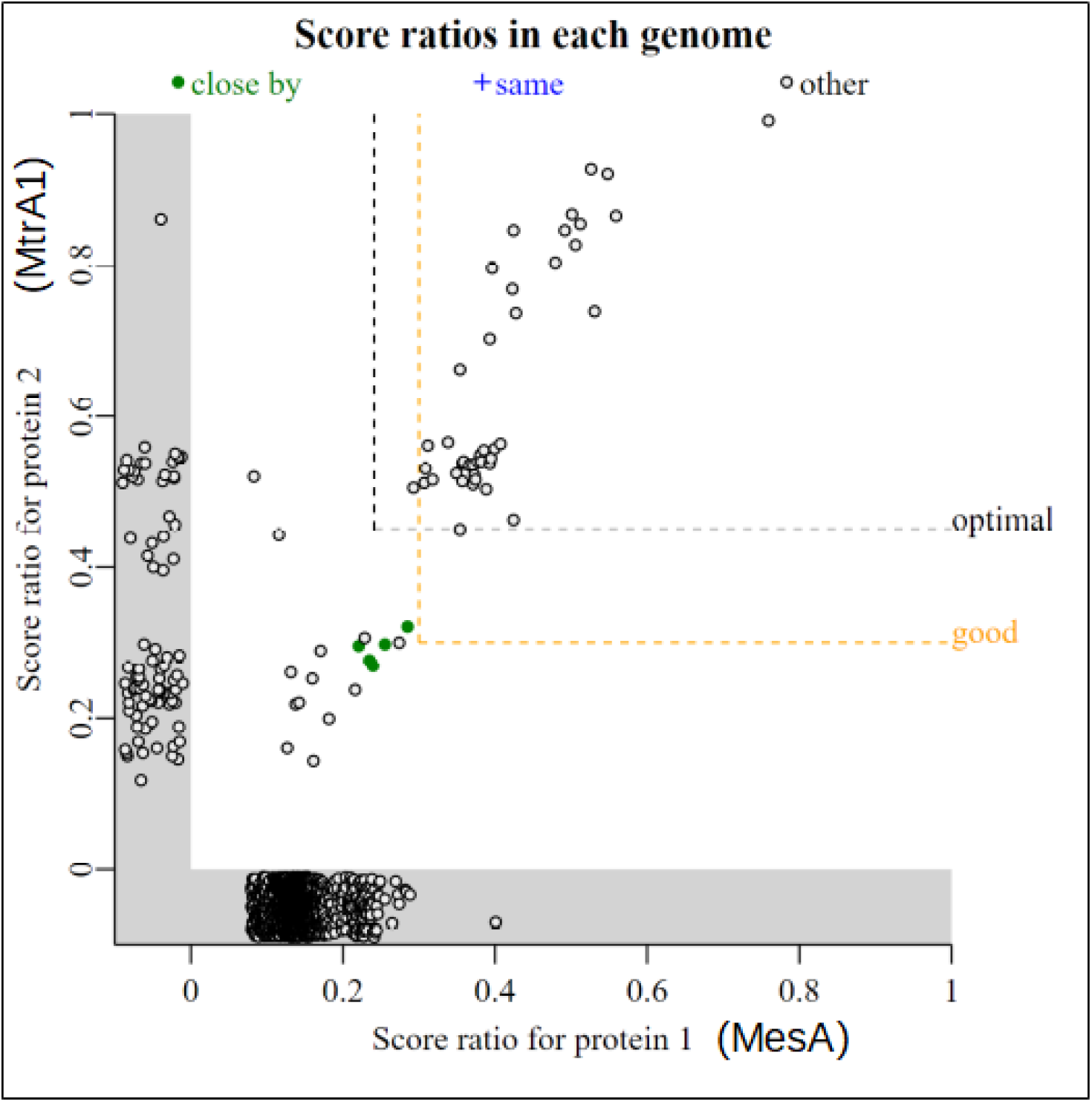
Comparing the presence/absence of two gene families. In this screenshot from *fast.genomics*, each point is a genome. The bit score ratio is the alignment score for the best hit, divided by the highest possible alignment score; it is a measure of how similar the best hit (if any) in that genome is to the query. The *x* axis shows the similarity to MesA from *M. marburgensis.* The *y* axis shows the similarity to MtrA1 from *M. marburgensis*. Genomes that have one protein, but not the other, are shown in the gray zones below zero. The labels for MesA and MtrA1 were added by hand.

*Fast.genomics* can go from a protein sequence to homologs from diverse bacteria and archaea in a few seconds. By default, it uses a database of 6,377 representative genomes, each from a different genus of bacteria or archaea. Alternatively, it can search within a larger set of genomes (up to 10 representatives per species) within any given taxonomic order. Either way, once it has computed a list of homologs, it can show gene neighborhoods (as in Figure 2B), compare presence/absence (as in Figure 6), or show the taxonomic distribution of a gene or a pair of genes.

We believe that *fast.genomics* is usually superior to other fast tools for these comparative genomics analyses because *fast.genomics* compares the query to all genes in the database on the fly (using mmseqs2, (Mirdita et al. 2019)). The other fast tools that we are aware of rely on ortholog groups, or pre-computed groups of similar proteins that ideally have the same function. In practice, ortholog groups are often either too broad -- they mix together proteins with different functions -- or too narrow, so that similar proteins that probably have the same function are missing from the group (Price and Arkin 2024). Either way, this can lead to less accurate or confusing results. However, *fast.genomics* may not identify remote homologs (roughly, under 30% amino acid identity). For the comparative genomics of broad protein families, such as PFams, we recommend GeCoViz for gene neighborhood analysis (Botas et al. 2022) and AnnoTree for co-occurrence analysis (Mendler et al. 2019). Finally, *fast.genomics* can only compare the presence/absence of two specified proteins. Given a protein family of interest, you can search for other protein families that have a similar occurrence across genomes using PhyloCorrelate (Tremblay et al. 2021).

### Finding candidates for a function with Curated BLAST for genomes

All the analyses so far began with a protein of interest. But often we start with a question about a specific organism, such as, does it have perchlorate reductase? Can it synthesize leucine, or break down phenylacetate? Or maybe we know that it can grow on phenylacetate, and we’d like to identify the pathway.

One way to look for a given protein function is to search the automated protein function annotations. However, a protein’s annotation is often non-specific, even if the protein is similar to a protein that is known to have that function, because of uncertainty as to which of several related functions the protein has.

Instead, Curated BLAST for genomes makes a list of characterized proteins whose annotations match the query, such as “perchlorate” or an Enzyme Commission (EC) number (Price and Arkin 2019). (The characterized proteins are taken from PaperBLAST’s database, see Table 2.) Then, it finds all homologs of these proteins in the genome of interest. Candidates can be checked further using PaperBLAST and other tools discussed above.

Also, sometimes proteins are missing from the genome annotation, either due to a misprediction of the open reading frame or due to a frameshift error in the genome sequence. After comparing the matching characterized proteins to the annotated proteins, Curated BLAST for genomes searches the six-frame translation of the genome. This will occasionally find proteins that were not annotated. If there is a frameshift, it is difficult to know if it is a true frameshift that renders the gene non-functional (a pseudogene), an error in the sequence, or (less likely) a protein that split into two pieces that still function. In our experience, frameshift errors are common in genomes that were sequenced solely using long reads (Pacific Biosciences or Oxford Nanopore).

### Tools for annotating pathways

Searching for individual protein functions can be cumbersome if there are many steps in the pathway of interest, and especially if there are multiple alternate pathways. For amino acid biosynthesis and the catabolism of small carbon sources, GapMind annotates the known pathways from bacteria and archaea (Price et al. 2020; Price et al. 2022). GapMind does not predict if the capability is present or not; rather, it compares the predicted proteins in the genome to all characterized proteins that carry out relevant enzymatic reactions or transport steps, and reports the best-supported path (Figure 7). Each step is color-coded by whether there is a high-confidence candidate. (Roughly speaking, a high-confidence candidate is at least 40% identical to an experimentally-characterized protein that has the function, and is less similar to characterized proteins that have other functions.) GapMind is a web-based tool that takes 10-40 seconds per genome, so it is convenient to run when needed.

**Figure 7:**
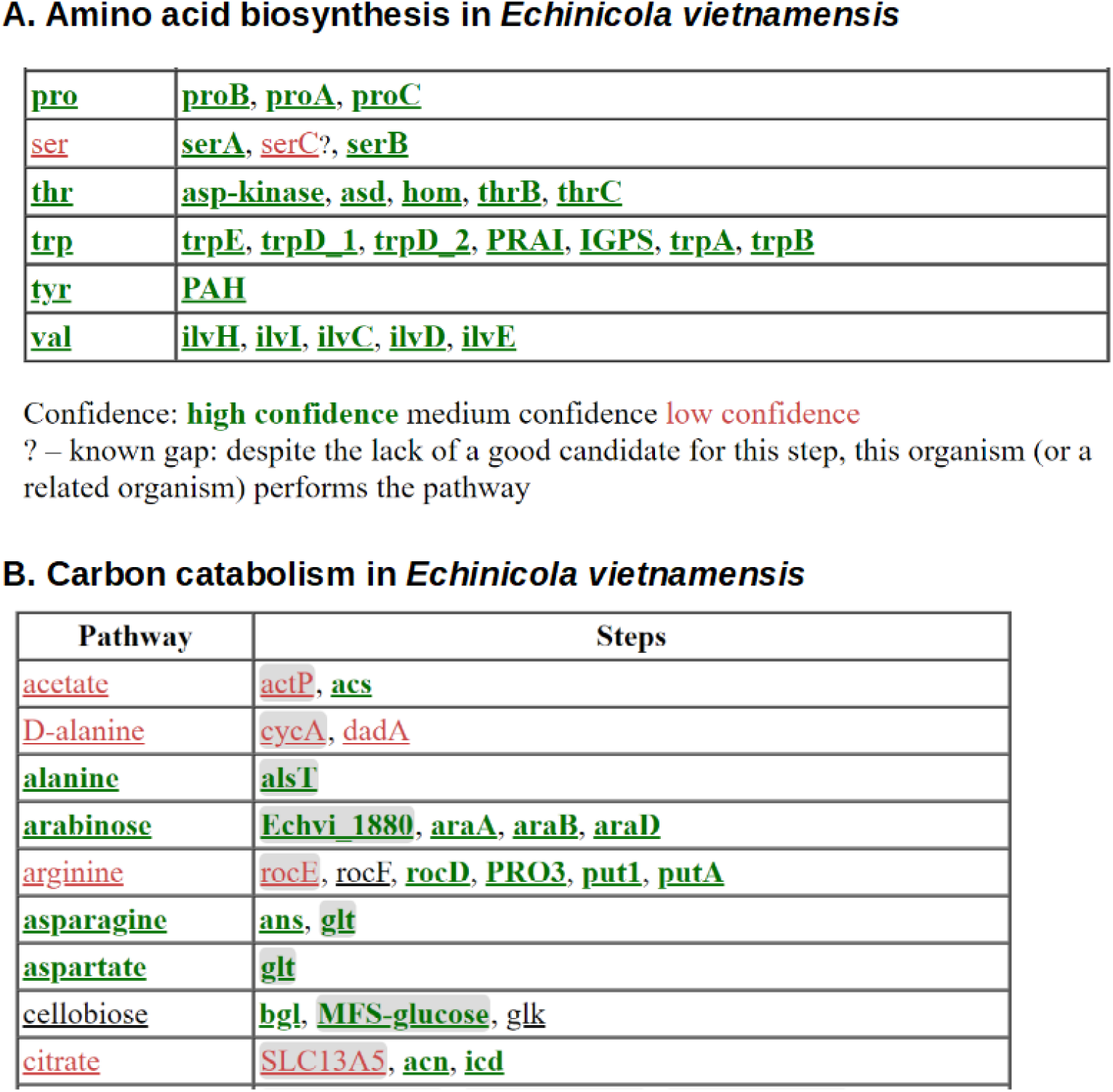
Annotating metabolic pathways with GapMind. (A) Amino biosynthesis in *Echinicola vietnamensis* DSM 17526. This screenshot shows biosynthetic pathways for five of the amino acids along with the key for the color-coding. (B) Carbon catabolism in *E. vietnamensis.* This screenshot shows catabolic pathways (or the lack thereof) for nine compounds. Transporters, which are more challenging to annotate, are shown with a gray background.

There are many other tools with similar goals that cover a different range of pathways than GapMind does. In particular, DRAM predicts the presence of metabolic pathways, mostly relating to energy production and central metabolism (Shaffer et al. 2020).

Regardless of which pathway annotation tool you use, the results are often ambiguous. For instance, what does it mean if the first and third steps in serine synthesis are present, but there’s no apparent protein for the second step (Figure 7A)? In *E. coli*, disrupting any one of the three enzymes results in a requirement for serine for growth, so the missing enzyme is definitely necessary. We refer to these missing steps -- which may well be encoded in the genome -- as gaps. In some cases, we can infer that the missing step is indeed present: if the other two steps are confidently annotated, and they are not known to be part of a different pathway, then there’s no other reason for the two steps to be present. (This logic is stronger if multiple species or genera have the same gap; otherwise, it is possible that the third gene was lost recently and the other two genes, although now useless, have not been lost yet.) Also, GapMind for amino acid biosynthesis identifies “known gaps.” If a related bacterium (from the same family) is known to synthesize the amino acid, even though the gene for the step cannot be found, then the absence of the step should not be viewed as evidence that the pathway is absent. In Figure 7A, the missing *serC* is a known gap because *E. vietnamensis* grows in a defined minimal medium without any amino acids (Price et al. 2020). SerC was probably replaced by a non-homologous or distantly-related protein that has not been identified yet. More broadly, analyses of gene fitness or gene neighborhoods, as described above, can sometimes identify the alternative proteins and fill the gaps (i.e., (Price et al. 2018a; Ashniev et al. 2022; Price et al. 2022)).

If an organism has a metabolic capability, then at least some of the genes for that capability can usually be identified, but not always. As an extreme example, we have not been able to determine how most Desulfovibrionales make L-serine, despite collecting large-scale genetic data from three representatives (Kuehl et al. 2014; Price et al. 2020; Trotter et al. 2023). These bacteria do not have a high-confidence assignment for any of the three steps of serine synthesis, yet they grow in a defined minimal medium,

If the above tools aren’t relevant to your question, we recommend using MetaCyc (Caspi et al. 2020) to browse the known metabolic pathways, followed by using Curated BLAST for Genomes to find candidates. Unfortunately, MetaCyc is no longer freely available; if you don’t have access, KEGG is an alternative (Kanehisa et al. 2016b).

Another way to annotate pathways is to use a genome-scale metabolic model. These models can work well if the model is curated to match the known physiology of the organism, but curation is laborious. Metabolic models that are generated entirely automatically are probably not accurate enough to be useful: in particular, the resulting predictions for amino acid auxotrophies or for carbon source utilization are not at all accurate (Gralka et al. 2023; Price 2023). In our view, given the challenges of automated annotation discussed above, we should expect automatically-generated models to be riddled with errors. In principle, automated gap-filling could be used to correct some of these errors, but in practice, automated gap-filling may introduce additional errors (Karp et al. 2018).

As an alternative to predicting capabilities from the genome, we can also consider the known capabilities of its relatives. For strongly conserved traits, such as many modes of energy production, the taxonomic classification of the organism could be as informative as the genome sequence. For instance, virtually all cyanobacteria can fix carbon dioxide. If a genome of a cyanobacterium appears to be missing a key gene for this pathway, it probably indicates an error in the genome or in the identification of open reading frames. Unfortunately, we do not know of a convenient web-based tool to predict traits from taxonomy; we usually rely on literature searches.

### The future

We expect the volume of high-throughput genetic data to increase dramatically. So far, Fitness BLAST finds reasonably-close homologs (≥40% identity) with specific phenotypes or cofitness for about a quarter of all bacterial proteins. We hope to reach the point where useful genetic data is available for homologs of most bacterial and archaeal proteins, but this will require far more data as well as new genetic tools so that randomly-barcoded transposon libraries become available for a greater diversity of bacteria and archaea.

We also expect that databases of the metabolic capabilities of bacteria will grow dramatically. Besides enabling more accurate predictions of metabolic capabilities from genome sequences, this will enable the identification of gaps in inferred pathways on a much larger scale. It may also make it easier to fill gaps by comparing the phylogenetic distributions of candidate proteins to those of the gaps.

We hope that the databases of experimentally-characterized proteins will improve, to cover more of the knowledge that is not currently available to automated tools. Unfortunately, curation is expensive: once a funding agency has spent ∼$100,000 for scientists to characterize a protein, they are not willing to spend ∼$300 for a curator to enter this information into a database (Karp 2016). In our view, this is penny-wise and pound-foolish. In any case, as automated text processing improves, curation might become partly automated, and costs should drop. Focusing curator effort on proteins with many homologs, but without close homologs that are characterized according to the curated databases, might also be a way to reduce costs. Alternatively, scientists could curate the functions of proteins when they publish papers about them (discussed in (de Crécy-Lagard et al. 2022)). Also, more papers could be linked to sequences via large-scale analysis of primer sequences, which are most often found in supplementary material (Haeussler et al. 2011).

Recently, high-quality structure predictions have become available for most proteins (Varadi et al. 2022). In principle, these structures could reveal the function of these proteins. So far, we have found structure prediction most useful for testing putative protein complexes and for finding remote homologs. It is not yet clear how to use AlphaFold and related approaches to predict the substrates of enzymes. Assuming that docking improves, but that accurate substrate prediction remains impossible, one strategy that may become broadly applicable is to combine docking with the gene neighbor method, also known as “pathway docking” (Zhao et al. 2013; Kumar et al. 2014; Calhoun et al. 2018).

## Materials and Methods

### Inferring protein functions from specific phenotypes

500 genes with specific phenotypes in carbon sources and nitrogen sources were taken from the December 2023 release of the Fitness Browser, and were checked manually. Many of them were quickly discarded because their phenotypes were detrimental, or their phenotypes did not show such a specific pattern when checked manually, or neither their annotations (from KEGG or from SEED (Overbeek et al. 2014)) nor their domain content (from PFam or TIGRFam (Haft et al. 2013), which are included in the Fitness Browser) suggested that they were enzymes or transporters. Among the remaining proteins, inferred functions for 96 proteins had previously been incorporated into the Fitness Browser’s reannotations; most of these were reported in the supplementary material of (Price et al. 2018b) or (Price et al. 2022). For another 90 proteins, we inferred functions using the Fitness Browser and the other interactive tools described above, especially PaperBLAST.

We double-checked the assigned functions for all proteins that we classified as annotation errors (by any of the four automated methods). This led to two ambiguous cases. First, the putative transporter PfGW456L13_4770 from *P. fluorescens* GW456-L13 is clearly important for L-asparagine utilization, while Swiss-Prot best hit, KEGG, and RefSeq annotate it as a glutamate/aspartate transporter. Since the main asparaginase appears to be cytoplasmic (PfGW456L13_740), L-aspartate transport cannot explain the phenotype of PfGW456L13_4770. But, mutants of PfGW456L13_4770 may also have a subtle defect during glutamate utilization (fitness = −0.9; we usually ignore effects with |fitness| < 1). Since the data suggested that asparagine is the primary substrate, this was still classified as an error for the automated methods. Second, the putative transporter AO356_14110 from *P. fluorescens* FW300-N2C3 is often annotated as “RarD” or as “chloramphenicol-sensitive protein RarD” because it is 48% identical to *E.coli* RarD. The molecular function on RarD is not known. Another member of this family (PA14_19160, 31% identical to AO356_14110) was proposed to be a citrate transporter, as it is genetically redundant with another citrate transporter during growth on citrate (Underhill and Cabeen 2022). AO356_14110 is specifically important for growth with D-serine as the nitrogen source (fitness = −1.5). Also, a close homolog is specifically important for growth with D-serine as either the carbon source or the nitrogen source (PfGW456L13_1142, 93% identity, fitness = −3.3 to −4.4.) These mutant phenotypes suggest AO356_14110 and PfGW456L13_1142 are involved in D-serine uptake, rather than chloramphenicol resistance. On the other hand, both of these genomes encode dsdX-type D-serine permeases, which are also important for utilizing D-serine. Our preferred explanation is that the rarD-type permeases are partially genetically redundant with dsdX-type permeases, but it is impossible to be sure. In any case, since chloramphenicol resistance seems unlikely to be the main role of these rarD homologs, we still classified those automated annotations as errors.

### Automated annotations

Annotations were derived from Swiss-Prot by using the best hit with E ≤ 10^-5^ and 70% coverage (both ways), as identified using NCBI protein BLAST+ against Swiss-Prot release 2023_05 (downloaded on January 12, 2024). KEGG-based annotations were from ghostkoala, run against prokaryotes + eukaryotes + viruses in January 2024. RefSeq annotations were from the corresponding genomes in RefSeq, downloaded in January 2024. Three of the 186 catabolic proteins are from genomes that were not in RefSeq, so we used RefSeq’s annotations of close homologs instead.

When determining the substrates implied by the best hit in Swiss-Prot, we considered the description and also the “Function” field. For instance, the protein PGA1_c29700 is usually annotated as a glycolate dehydrogenase, based on its similarity to the *E. coli* GlcF protein. In the fitness data, PGA1_c29700 is important for the utilization of D,L-lactate or D-lactate (but not L-lactate), which suggests that it is a D-lactate dehydrogenase. The Swiss-Prot description for the best hit (which is *E. coli* GlcF) states that GlcF has similar activity with D-lactate as with glycolate, so activity on D-lactate should be expected; thus, the best-hit annotation from Swiss-Prot was considered correct. Similarly, for CLEAN, which reports only the Enzyme Classification (EC) number, we consulted the description of the EC number. In this case, CLEAN assigned the EC number 1.1.99.14, whose description reports that it also acts on D-lactate. Because users of protein annotations usually consider only the textual description, we may have slightly overstated the accuracy of Swiss-Prot best hits and CLEAN. In contrast, for KEGG and RefSeq, we did not look at further information besides the text that was provided by ghostkoala or by RefSeq itself.

### PaperBLAST and related databases

Since the original publication describing PaperBLAST (Price and Arkin 2017), we have incorporated additional resources into its database for linking proteins to papers. All of the resources are listed in Table 3. The additional resources are:

- BioLiP is a database of biologically relevant ligands in protein structures (Zhang et al. 2024). All of these ligand-binding proteins are included in PaperBLAST’s database. Even if there is no link to a publication, the fact that the protein bound the ligand can be informative.
- MetaCyc is a database of metabolism (Caspi et al. 2020). Only proteins (or complexes) that link to both a sequence and to one or more papers are included in PaperBLAST’s database.
- PaperBLAST incorporates a small subset of the European Nucleotide Archive (ENA, (Cummins et al. 2022)) with experimental evidence. PaperBLAST scans nucleotide entries from the “STD” class (roughly, small-scale sequencing projects) for coding sequences (CDS features) with the /experiment tag. The corresponding proteins are included in PaperBLAST’s database if the nucleotide entry links to one or more papers in PubMed. Also, to filter out genes whose transcription or translation was detected, but whose function might not have been not studied, entries with more than 25 such proteins, or papers that link to more than 25 proteins in this way, are excluded.
- The incorporation of the experimentally-characterized subset of TCDB, a database of transporters (Saier et al. 2016), was described previously (Price et al. 2022).
- RegPrecise is a database of predicted regulons, as reconstructed by comparative genomics (Novichkov et al. 2013). Because RegPrecise often includes predictions for several similar transcription factors from related species, we clustered the predicted regulators at 70% identity and 80% coverage with usearch (Edgar 2010). One representative of each cluster is included in PaperBLAST’s database.
- The Fitness Browser reannotations are a collection of proteins whose function was inferred from RB-TnSeq data (mostly from (Price et al. 2018b; Price et al. 2022)).
- PRODORIC is a database of experimentally-characterized regulons (Dudek and Jahn 2022). The regulators that have UniProt identifiers were incorporated into PaperBLAST.

The database of characterized proteins (Table 2) is a subset of the full PaperBLAST database. Heuristics for removing uncharacterized proteins were described previously (Price et al. 2020).

SitesBLAST uses a separate database, based on BioLiP and on Swiss-Prot entries whose sequence features have experimental evidence. To incorporate BioLiP into SitesBLAST, we cluster the protein sequences at 90% identity and 80% coverage with CD-HIT (Fu et al. 2012). From each cluster, we select representatives so that every ligand that binds any member of the cluster is included in a structure.

## Availability of tools and data

Except for MetaCyc and Google Colab, all of the tools that we discussed are freely available. Up-to-date databases for PaperBLAST, the Fitness Browser, and GapMind are available from their websites. The database for the February 2024 release of the Fitness Browser is archived at figshare (https://doi.org/10.6084/m9.figshare.25236931.v1). (This is identical to the version we used for identifying catabolic genes in bacteria, except that it also includes data from two archaea.) The database for the December 2023 release of PaperBLAST is archived at figshare (https://doi.org/10.6084/m9.figshare.25254562.v1). The 500 proteins with specific phenotypes that we examined and the 186 enzymes and transporters whose functions we identified, as well as the annotations from automated tools, are available at http://tinyurl.com/3dzn9f2b.

## Acknowledgements

This material by ENIGMA-Ecosystems and Networks Integrated with Genes and Molecular Assemblies (http://enigma.lbl.gov), a Science Focus Area Program at Lawrence Berkeley National Laboratory is based upon work supported by the U.S. Department of Energy, Office of Science, Office of Biological & Environmental Research under contract number DE-AC02-05CH11231.

